# Neural synchrony between prefrontal and visual cortex supports visual working memory

**DOI:** 10.64898/2026.06.05.730488

**Authors:** Mrugank Dake, Sangita Dandekar, Clayton E. Curtis

**Affiliations:** Department of Psychology, New York University, New York, NY 10003; Center for Neural Science, New York University, New York, NY 10003; Department of Psychological and Brain Sciences, Dartmouth College, Hanover, NH 03755

**Keywords:** Working memory, frontovisual synchrony, beta band oscillations, spatial memory

## Abstract

Working memory appears to depend on neural mechanisms that are distributed across the brain. Specifically, neural activity persists in the prefrontal cortex while memories are maintained, and at the same time, the visual contents of memory can be precisely decoded from the patterns of activity in visual cortex. Contemporary models attempt to account for these findings by positing that higher-order areas, like prefrontal cortex, somehow control memory storage by recruiting encoding mechanisms in sensory cortices. Demonstrating how prefrontal cortex influences working memory representations in visual cortex is methodologically challenging and direct evidence remains scarce. Here, we leveraged the excellent temporal resolution of magnetoencephalography to test hypotheses about how synchronization of neural activity between prefrontal and visual cortex coordinates working memory representations. During a visuospatial working memory task in humans (both sexes), increased power in 𝛽-band activity persisted throughout memory maintenance, and changes in its topography over visual cortex predicted both memorized locations and trialwise memory errors. Moreover, neural activity in the 𝛽-band synchronized between the prefrontal and visual cortices during memory. Not only do these findings align with a large body of work demonstrating that working memories are widely distributed across the brain, but they also help explain how the prefrontal and sensory cortices communicate during memory.

**New & Noteworthy:** While canonical models emphasize prefrontal memory storage, recent theories propose distributed interactions between executive control and sensory areas. Using magnetoencephalography, we demonstrate that 𝛽-band oscillations over visual cortex track specific memory content and synchronize with prefrontal cortex activity during maintenance. Crucially, this localized sensory activity directly predicts trial-by-trial human memory errors. These findings provide critical empirical evidence for sensory recruitment models, revealing a dynamic, oscillatory mechanism coordinating working memory.

## Introduction

Working memory, the ability to store information that can guide behavior, is an essential ability that higher cognitive functions depend upon (Baddeley, 2003), and cognitive symptoms found in a number of psychiatric and neurological diseases, especially schizophrenia, may stem from working memory dysfunction (Snitz et al., 1999). Therefore, clinical applications critically depend on a better understanding of the neural mechanisms of working memory (Millan et al., 2012). Our best evidence is that neuronal activity in the macaque dorsolateral prefrontal cortex persists during the retention intervals of working memory tasks (Fuster and Alexander, 1971; Bruce et al., 1985; Funahashi et al., 1989; Miller et al., 1996). Such persistent activity provides a candidate neural mechanism for working memory storage and is the cornerstone of canonical theories of working memory (Compte et al., 2000; Wang, 2001). Accordingly, sustained activity among prefrontal neurons selective for the memorized feature stores the feature until the memory is later used to guide behavior (Riley and Constantinidis, 2015).

An accumulation of more recent evidence suggests that such theories may be too narrowly focused both in terms of anatomy (i.e., prefrontal cortex) and function (i.e., memory storage). Indeed, persistent activity has been observed in many parts of the cortex and even subcortex (Miller and Desimone, 1994; Pesaran et al., 2002; Bisley et al., 2004; Funahashi, 2013; Hayden and Gallant, 2013; Rossi-Pool et al., 2017; Bastos et al., 2018; Sadeh et al., 2018), suggesting that memory representations are widely distributed beyond just the prefrontal cortex (Christophel et al., 2017). Indeed, non-invasive electrophysiological studies in humans using memory-guided saccade paradigms have long demonstrated that posterior parietal and occipital areas track spatial memory metrics over delays <u>(Medendorp et al. 2007; Van Der Werf et al. 2008)</u>. Moreover, the multivariate patterns of neural activity in early visual cortex, including primary visual area, V1, can be used to decode the contents of visual working memory (Harrison and Tong, 2009; Rahmati et al., 2018). While precise decoding has been replicated many times now, it remains perplexing because persistent activity, the neural mechanism that presumably stores working memory representations, is often absent in these early sensory areas (Fuster, 1990; Chelazzi et al., 1998, 2001; Zaksas and Pasternak, 2006; Mendoza-Halliday et al., 2014; Leavitt et al., 2017); but see (Supèr et al., 2001; van Kerkoerle et al., 2017).

While debates continue about the relative importance of prefrontal and sensory cortices for working memory (Curtis and D’Esposito, 2003; Pasternak and Greenlee, 2005; Sreenivasan et al., 2014; Curtis and Sprague, 2021), new theories propose that working memory depends on critical interactions between these areas. For instance, feedback signals from the prefrontal cortex may somehow recruit the encoding mechanisms in sensory cortex for memory storage (Curtis and D’Esposito, 2003; Postle, 2006; Serences, 2016; Comeaux et al., 2023). In these models, the prefrontal cortex’s role in working memory is reframed more in terms of executive control rather than storage. On the one hand, such a reframing aligns well with a host of experimental findings, including poor stimulus selectivity of prefrontal neurons (Miller et al., 1996; Pasternak and Greenlee, 2005; Mendoza-Halliday and Martinez-Trujillo, 2017), inability of prefrontal areas to decode memory content (Riggall and Postle, 2012; Emrich et al., 2013; Curtis and Sprague, 2021), and that lesions do not impact working memory performance in humans (D’Esposito and Postle, 1999; Mackey et al., 2016a, 2016b) and only inconsistently do so in macaques (Rushworth et al., 1997). On the other hand, evidence for the proposed interaction and the mechanisms by which feedback from prefrontal cortex recruits sensory areas remains poorly understood.

Here, we leveraged the temporal resolution of magnetoencephalography (MEG) to test if feedback from human prefrontal cortex synchronizes neural activity within the visual cortex as a means to recruit the relevant, memory-specific neural populations. In brief, we found that power changes in the 𝛽-band not only predicted working memory content and performance, but also neural activity in the 𝛽-band synchronized between prefrontal and visual areas during the retention interval.

## Methods

### Participants

Twenty-eight neurologically healthy participants (12 males, 16 females; all right-handed, mean age 25.3 (range: 18 - 43 years) were recruited to participate in a memory-guided saccade (MGS) task inside an MEG scanner. All participants reported normal or corrected-to-normal vision and no previous history of neurological disorders. Participants provided written informed consent and all procedures were pre-approved by the Institutional Review Board (IRB) at New York University (NYU). One participant had noisy neural data and six participants had degraded gaze data and were eliminated from all analyses. Finally, we have twenty-one participants in the study. Participants were compensated $30/hr for their time.

### Memory-Guided Saccade (MGS) Task

The experiment was controlled with MGL toolbox (https://gru.stanford.edu/doku.php/mgl/overview) and projected (InFocus LP425) into the magnetically shielded room (Vacuumschmelze, Hanau, Germany) onto a screen placed 57cm away from the participants. Participants performed an MGS task **(Figure 1A)**. Each trial began with a white fixation dot presented at the center of the screen for 1000ms, followed by a stimulus (a gray dot) presented at one of the 10 locations at a 9-degree eccentricity, which flashed for 200 ms. This was followed by a variable delay of either 1500ms or 3500ms, during which participants were instructed to fixate at the central white fixation dot. The disappearing fixation cross provided the go cue and the participants were instructed to make a saccade toward the location of the target in their memory within 1000ms. This was followed by the display of a green dot at the target location that served as feedback for 1500ms, during which participants were instructed to make a corrective saccade and fixate on the green dot until it disappeared. Finally, a gray dot at the center of the screen marked the end of the trial. The targets were presented at specific polar angles (0°, 25°, 50°, 130°, 155°, 180°, 205°, 235°, 310°, 335°). Each run had 35 trials. Participants performed 8-10 runs of the experiment.

**Figure 1:**
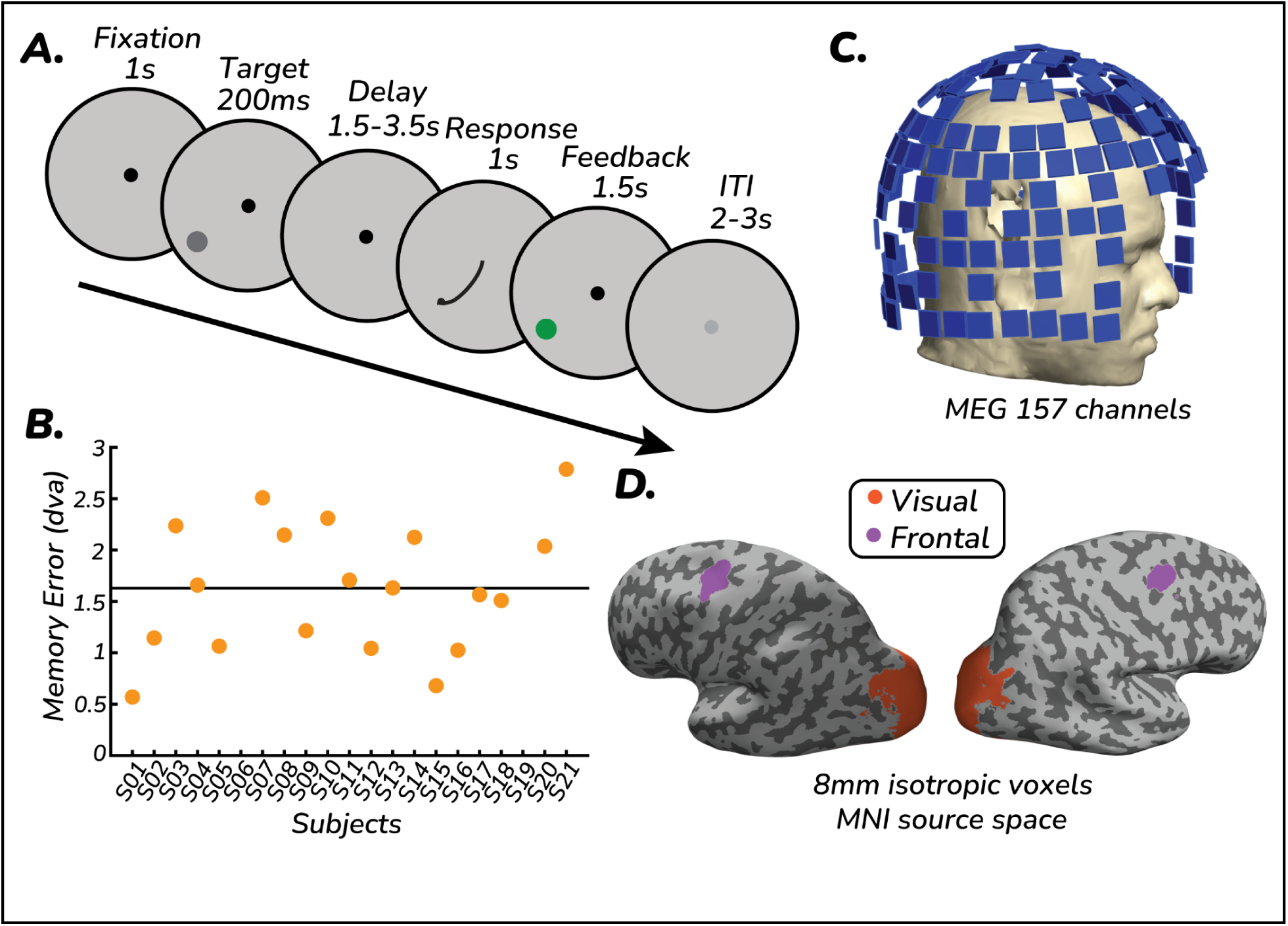
Experimental paradigm, behavioral performance and MEG source localization. **A.** Memory-guided saccade (MGS) task. Participants fixated on a central fixation point for 1s before a brief target (200 ms) was flashed at one of the 10 polar angle locations. Following a variable delay (1.5-3.5s), the disappearance of the fixation dot indicated a go cue, after which participants made a saccade to the remembered target location. Feedback was provided via a green dot at the actual target location. **B.** Mean saccade error (degrees of visual angle, dva) across participants (*N* = 21). The horizontal line indicates group mean error 1.58 +/- 0.15 dva, demonstrating high spatial accuracy. **C.** MEG recording. Data were recorded using a supine 157-channel axial gradiometer KIT system. The schematic illustrates the sensor layout relative to the standard digitized head model. Region of interest (ROI) definitions. Visual (orange) and Prefrontal (purple) ROIs projected onto MNI source space (8mm voxels). ROIs were defined using Wang atlas (Wang et al., 2015) projected on the FreeSurfer surface for visualization.

### Eye tracking

Gaze of the participants was recorded using an MEG-compatible fiber optic Eyelink 2K (Ontario, Canada). The right-eye gaze data were sampled at 1000Hz. For all participants, a 9-point calibration was performed prior to the onset of the recording session and repeated occasionally whenever a participant took a break or gaze data drifted during a run. Participants were instructed to refrain from blinking and breaking fixation during the trial until the response onset. The gaze data were preprocessed offline using iEye (https://github.com/clayspacelab/iEye), an in-house gaze preprocessing and analysis toolbox written in MATLAB. Additionally, the gaze data were calibrated to the targets offline. The data were first preprocessed to remove blinks, which were identified as pupil diameter under 1 percentile for each recording, and the gaze data 100 ms pre and post-blink onset were removed. The gaze data were then smoothed using a Gaussian smoothing kernel to estimate velocity. Saccades were detected as gaze transitions with velocities greater than 30 deg/s, having an amplitude greater than 0.25 degrees, and a duration greater than 7.5 ms. The gaze data were also corrected for instrumental drift by demeaning the gaze data during fixation and delay epochs. Lastly, we performed an offline calibration using fixation during corrective saccades as a proxy for each trial to account for any instrumental or head-movement-related drift.

### Magnetoencephalography (MEG) recording and preprocessing

Neural data were recorded using a 157-channel axial gradiometer KIT system (Kanazawa Institute of Technology, Kanazawa, Japan) **(Figure 1C)**. The data were sampled at 500Hz. Participant head location was tracked using the head position locator coils. MEG data is prone to environmental noise and we removed the non-periodic low-frequency noise using CALM (Continuously Adjusted Least Squares Method) noise reduction (Adachi et al., 2001) in MEG160 (Yokogawa Electric Corporation and Eagle Technology Corporation, Tokyo, Japan). MEG data were preprocessed using Fieldtrip (Oostenveld et al., 2011). Specifically, the data were filtered using a sixth-order high-pass Butterworth filter at 0.5 Hz. The data were then manually screened for artifacts and noisy or flat recording channels. The flagged recording channels were removed and interpolated using weighted interpolation with neighboring channels. Independent Component Analysis (ICA) was performed using the ‘*runica*’ algorithm in Fieldtrip on this minimally preprocessed data, with the number of components extracted determined by the rank of the data. The components were then screened for blinks, eye movements, iron-cross (a signature of MEG recording at NYU, likely associated with NYC subway noise), and heartbeats. The data were then converted into planar gradients (Bastiaansen and Knösche, 2000). All further time-frequency and decoding analyses were performed on these planar gradients.

### Source Space Analyses: Forward Model Generation

Prior to the MEG scan, participants’ head shapes were digitized using a Polhemus FastSCAN-II. Since individual high-resolution T1-weighted anatomical MRI images were not available, we coregistered Montreal Neurological Institute (MNI) brain to individual head shapes. This was done using a two-step procedure. The first step involved minimizing the procrustes distance between MNI fiducials and native fiducials:

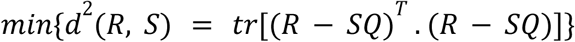

where, *R* is the matrix of reference fiducials (e.g., MNI coordinates), *S* is the matrix of native subject fiducials, *Q* is an optimal orthogonal rotation matrix that aligns *S* to *R*, and *tr* is the trace of the matrix.

Next, these semi-aligned MNI brains were then manually aligned to closely match the native participant fiducials and head shape. The obtained transformation matrix was then used to align the MNI singleshell head model to individual head shapes.

### Source Space Analyses: Inverse Model

The preprocessed MEG sensor data were epoched, lowpass filtered at 55Hz, and downsampled at 120Hz for computational efficiency. The covariance matrix was estimated using all trials, and the downsampled MEG data were then projected into the subject-aligned MNI source space with isotropic 8mm voxels using Linearly-Constrained Minimum Variance (LCMV) beamformer (Van Veen et al., 1997) with regions of interest (ROI) estimated from Wang atlas (Wang et al., 2015) **(Figure 1D)**. The source space data were then bandpass filtered in 𝛽-band (15-25 Hz), and the instantaneous power and phase were estimated using the Hilbert transform.

### Spectral Analyses

The preprocessed MEG sensor data, projected into the subject-aligned MNI source space, was further analyzed in the frequency domain. To estimate time-varying power for decoding analyses, the source space data for each trial were bandpass filtered using a fourth-order Butterworth filter. The analytical signal was then computed via the Hilbert transform, and the instantaneous power was derived as the squared magnitude of the analytic signal. Power estimates were subsequently baseline-corrected, defined as the relative change from the average power across all trials at each source. Finally, the time-varying power estimates were smoothed and downsampled to a 25-ms temporal resolution utilizing a 100 ms sliding averaging window (±50 ms from the center timepoint).

### Relative Power

To visualize the topography of 𝛽-band power, we computed relative power at each spatial location. The relative power at each location was defined as:

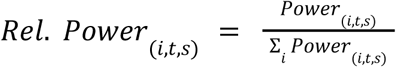

where, *Rel. Power_(i,t,s)_* is the relative power at spatial location *i*, time-point *t,* and source *s*. *Power_(i,t,s)_* is the baseline-corrected power, and the denominator is the sum of baseline-corrected power over all spatial locations.

### Support Vector Regression

Stimulus location was decoded from neural activity using a support vector regression (SVR) model with a radial basis function (RBF) kernel. We restricted our decoding analyses to visual and prefrontal ROIs designated by the Wang probabilistic atlas (Wang et al., 2015). Prior to decoding, trials containing invalid or missing eye-tracking data (e.g., undetermined initial saccade landing point) were excluded. For each source within the specified ROI, the instantaneous power was *z*-scored across all valid trials at each target location and time point.

Since the stimulus location varied continuously by polar angle, we trained two independent SVR models to predict the sine and cosine components of the polar angle:

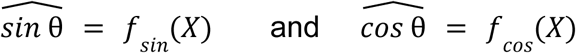

where, *X* represents the spatial pattern of normalized, baseline-corrected power in 𝛽-band across all sources within the ROI at a given time point.

The estimated stimulus angle was then reconstructed from the predicted sine and cosine components:

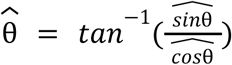

For each participant, we trained and evaluated a model at each time point using a leave-one-out cross-validation (LOOCV) procedure. This approach yielded a predicted stimulus angle based exclusively on 𝛽-band activity for each trial and time point. The SVR prediction error was defined as the minimum circular distance (angular error) between the reconstructed angle and the true angle.

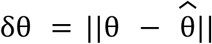

For behavioral correlation with decoding, we utilized these single-trial predictions to categorize trials into quantiles based on memory performance (initial saccade error).

### Coherence Analyses

To characterize task-related functional connectivity between prefrontal and visual regions, we performed seed-based connectivity analyses in source space. We utilized complex source-space data corresponding to 𝛽-band (15-25 Hz). A centered sliding time window of 400ms was applied at each time point to compute cross-spectral density.

To circumvent spurious phase-coupling estimates driven by signal leakage and volume conduction (Yoshinaga et al., 2020; Yu, 2020), we utilized the imaginary part of coherency (*ImCoh*). Functional connectivity was computed between predefined ROI pairs (left/right visual and left/right prefrontal regions) defined by the Wang atlas. Specifically, imaginary coherence was calculated as the absolute value of the imaginary component of the cross-spectrum normalized by the individual power spectra for the seed and target sources:

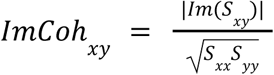

where, *S_xy_* is the cross-spectral density computed across all trials, and *S_xx_* and *S_yy_* are the corresponding auto-spectra for sources *x* and *y*.

To facilitate group-level comparisons and account for inherent differences in baseline connectivity across participants, the raw *ImCoh* values were *z*-scored relative to the mean and standard deviation of the pre-stimulation fixation period (-1 to 0s). The resulting *z*-scored time series was averaged across all vertices and within each ROI to produce a single connectivity metric per ROI pair for each participant.

### Statistical analyses

All statistical analyses were performed using custom routines implemented in Python. To compare task-modulated synchrony across conditions and ROI pairings, we focused on two primary contrasts: cross-hemispheric vs within-hemispheric pairs for the same prefrontal ROI, relative to the target stimulus location. For each contrast, we employed a non-parametric sign-flipping permutation test (*N* = 21). This approach constructs a null distribution by randomly inverting the sign of each participant’s difference score over 9999 iterations. The two-tailed *p*-value was defined as the proportion of permutations where the absolute mean difference exceeded the observed value. To address the temporal dimension of our coherence data, we identified significant temporal points (*p* < 0.05, not corrected for multiple comparisons). Effect sizes for key comparisons were reported using Cohen’s *d*. Significance thresholds were maintained at *p* < 0.05 (∗), *p* < 0.01 (∗∗).

## Results

### Behavioral Results

Memory errors were computed as the Euclidean distance between the initial saccade landing point and the true target location. Participants made an error of about 1.58 ± 0.15 degrees of visual angle (dva) **(Figure 1B)**. Additionally, we measured saccade reaction time as the time from response cue presentation to saccade onset. Participants made saccade responses within 356 ± 8 ms **(Supplementary Figure 1)**.

### 𝛽-power enhancement during working memory delay in visual sources

Power spectra were computed for visual and prefrontal ROIs in MNI space. We observed prominent peaks in α (8-12 Hz) and 𝛽 (15-25 Hz) frequency bands over the visual and prefrontal sources. Examining the differences in power during fixation and working memory delay, we observed an enhanced power in 𝛽 band **(Figure 2A)** during working memory delay in visual sources, and to a lesser extent in prefrontal sources **(Figure 2C)**. We also examined the time-frequency spectrogram, computing relative power throughout the trial with respect to fixation, across a range of frequencies (2-40 Hz). Both visual and prefrontal sources exhibited a broadband stimulus-locked activity profile, followed by a brief suppression of power in lower frequency bands and enhanced power in 𝛽-band that persisted throughout the working memory delay **(Figure 2B, D)**.

**Figure 2:**
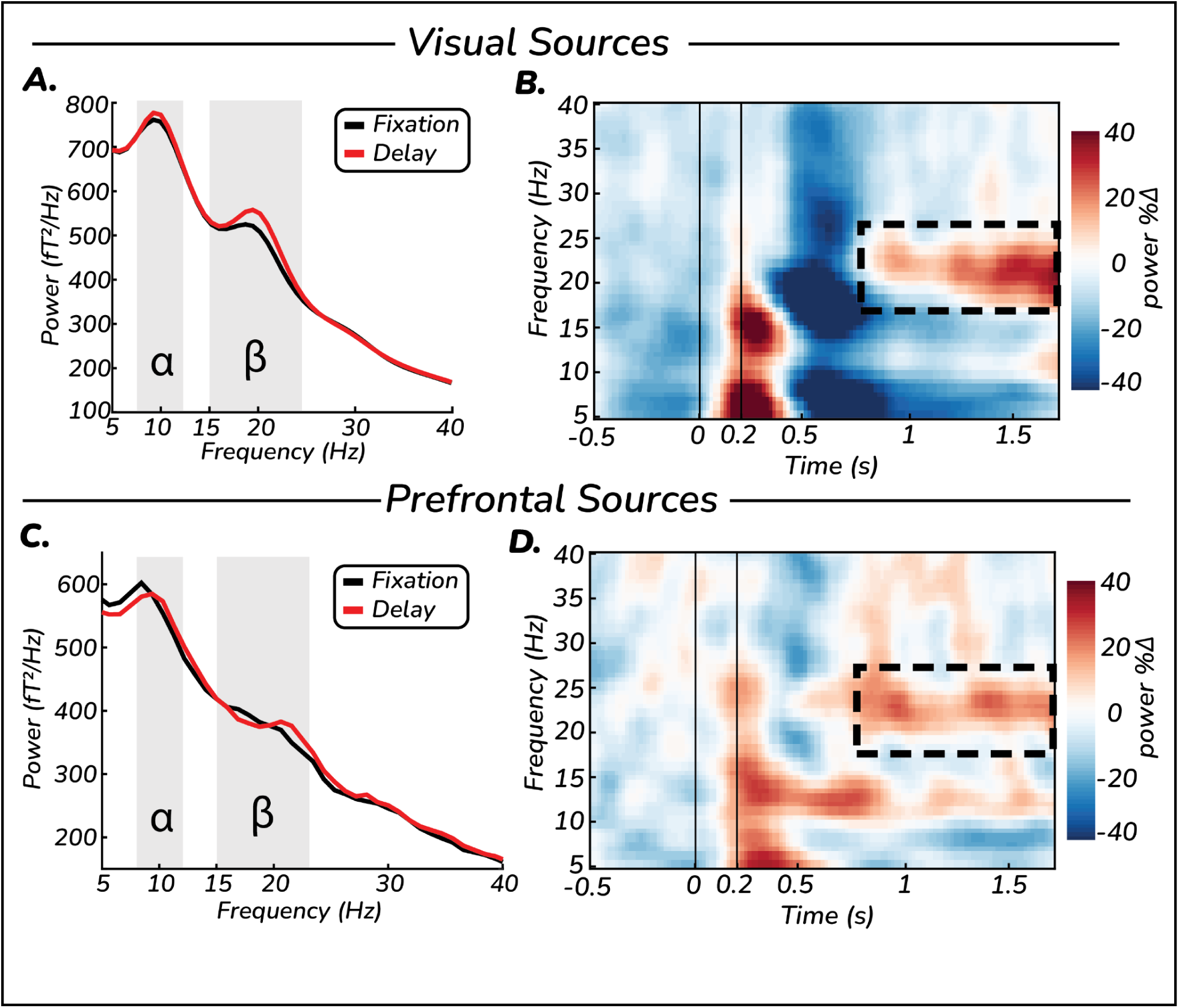
Spectral and temporal dynamics of neural activity during working memory delay. A, **C.** Power spectral density (PSD) for visual (A) and prefrontal (C) sources. Black curves represent the fixation epoch (-1 to 0s), and red curves represent the memory delay (averaged over 0.7-1.7s). Shaded gray regions denote the alpha (α: 8-12 Hz) and beta (𝛽: 15-25Hz) bands. Both regions exhibit distinct peaks in these bands, with a selective enhancement of 𝛽 power during working memory delay period compared to fixation. **B, D.** Time-frequency spectrograms for visual (B) and prefrontal (D) sources. Color maps represent the percentage change in power relative to the fixation baseline. Vertical lines indicate stimulus onset (0s) and offset (0.2s). Following an initial broadband stimulus-evoked response, both regions show a sustained increase in 𝛽 power (dashed black boxes) that persists throughout the maintenance interval.

### 𝛽-power tracks working memory content

We then investigated relative power in 𝛽-band across all sources during working memory delay by binning the trials into six groups of spatial locations. We observed a topographic change in the pattern of 𝛽-band power over visual cortex as a function of target polar angle **(Figure 3A)**. Specifically, for targets on the right hemifield, we observed an enhanced 𝛽 power over right visual sources and reduced 𝛽 power over the left visual sources. Similarly, for targets on the left hemifield, we observed an enhanced 𝛽 power over left visual sources and diminished 𝛽 power over the right visual sources. This is consistent with the well-established finding of hemispheric lateralization of low-frequency activity in the context of visuospatial working memory (Worden et al., 2000; Medendorp et al., 2007; Foster et al., 2017), likely reflecting an inhibitory gating mechanism where power increases in the task-irrelevant hemisphere to protect the memory trace from sensory interference (Jensen and Mazaheri, 2010).

**Figure 3:**
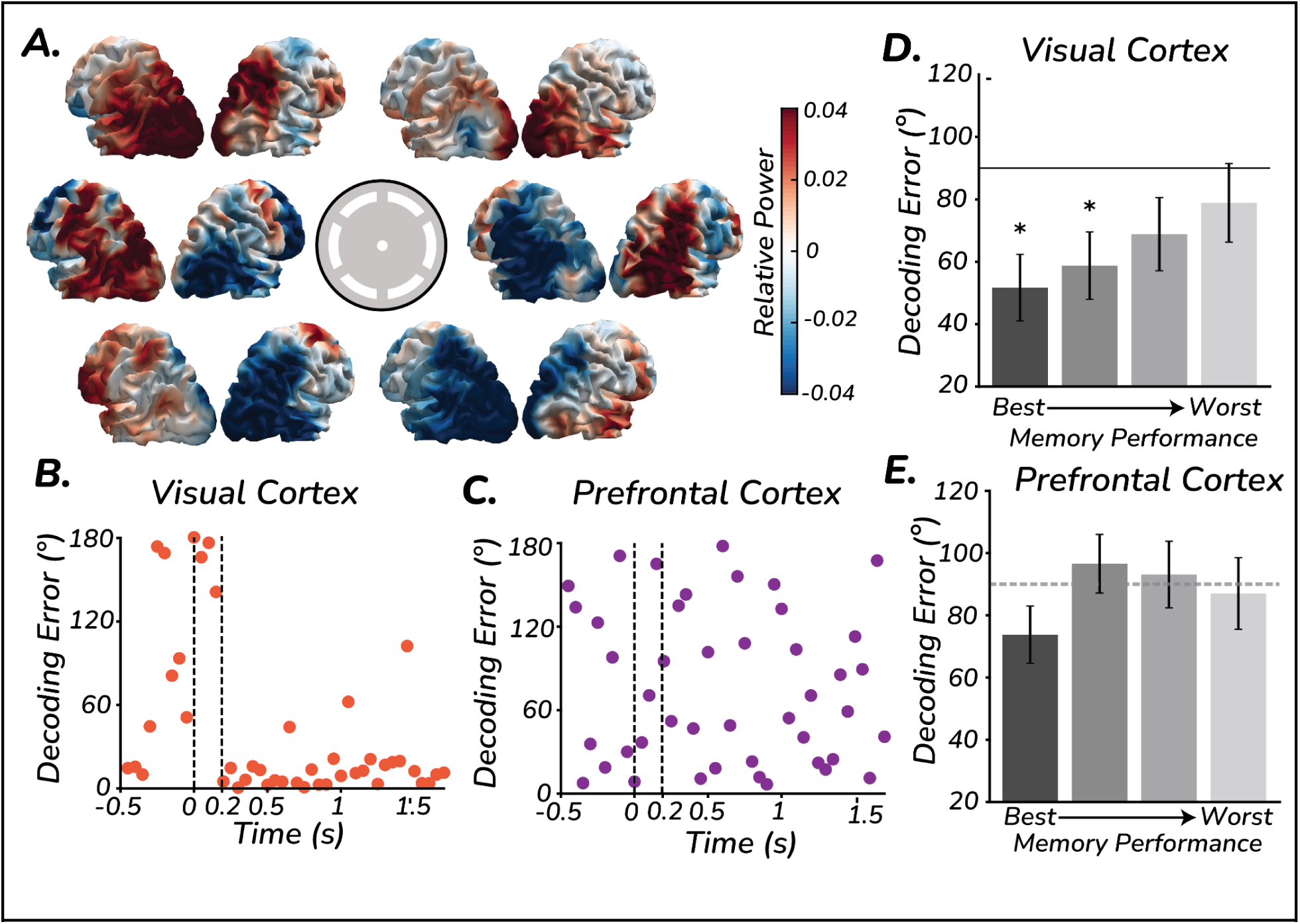
Visual, but not prefrontal, 𝛽**-band activity tracks high-fidelity working memory content. A.** Topography of 𝛽-band power over cortical sheet in MNI space during working memory delay, binned into six spatial target locations (wedges in the center). Relative power was calculated as the power at a specific target bin location relative to the mean power across all other bins. Note the characteristic spatial profile: enhanced 𝛽-band power in the ipsilateral hemisphere and suppressed power in the contralateral hemisphere relative to the target, and the rotational topography of this power as a function of target polar angle. **B, C.** Temporal dynamics of spatial decoding. Decoding error (degrees) for target location over time from 𝛽-band power in visual (B, orange) and prefrontal (C, purple) ROIs. Vertical dashed lines indicate stimulus onset and offset. High-fidelity working memory target location is reliably decoded from visual sources throughout the delay, whereas decoding from prefrontal sources remains at chance levels. **D, E.** Neural decoding predicts working memory performance. Trials were divided into four quantiles (Q1-Q4) based on memory error (best to worst, left to right). In visual cortex (D), decoding error was significantly different from chance for trials with best performance and not for worst, with a monotonous trend indicating correlation between behavioral performance and decoding precision. This relationship is notably absent in the prefrontal cortex (E). Error bars represent SEM; * denotes significant difference from chance (*p* < 0.05).

We further investigated stimulus-selectivity in the topography of 𝛽-power by training a Support Vector Regression (SVR) at each time point for visual and prefrontal sources separately using leave-one-out cross-validation (LOOCV). The model successfully decoded the target location from the topography of power during working memory delay in visual sources **(Figure 3B)**. However, this decoding was not significant in prefrontal sources **(Figure 3C)**. This is consistent with fMRI studies showing precise decoding of working memory contents from visual cortex (Harrison and Tong, 2009; Rahmati et al., 2018; Duan and Curtis, 2024) but not from prefrontal cortex (Christophel et al., 2012; Riggall and Postle, 2012).

In our task design, stimulus location and working memory content are spatially indistinguishable. To investigate whether 𝛽 power in visual cortex tracked stimulus location or working memory content, we divided each participant’s data into four quantiles based on their memory performance. We observed that the decodability of stimulus location from our decoder correlated with participants’ memory performance. Specifically, trials on which participants made the smallest memory errors, the decodability of the regressor was more precise compared to trials on which participants made large memory errors. Moreover, this trend was only observed in visual sources **(Figure 3D)** but not in prefrontal sources **(Figure 3E)**. This provides strong evidence for 𝛽-band activity in early visual cortex tracking working memory content.

It is noteworthy that ɑ-band power (8-12 Hz) also exhibited a stimulus-selective topography that allowed for significant spatial decoding **(Supplementary Figure 2)**. This activity, however, did not track memory performance in the same manner as 𝛽-band power. Specifically, ɑ-band decoding fidelity remained stable across all performance quantiles **(Supplementary Figure 2 D, E)**, suggesting a functional dissociation where 𝛽-band activity specifically reflects the fidelity of the stored representation, whereas ɑ-band activity may reflect a more genreal state of spatial orienting or suppression.

### Prefrontal cortex and visual cortex synchronize in 𝛽 band during working memory

To characterize the rhythmic coordination between the prefrontal and visual cortices during the task, we computed the imaginary coherence (*ImCoh*) in 𝛽 band. By focusing on the imaginary component of coherency, we isolated phase-synchronized interactions that are not attributable to instantaneous volume conduction. Our analysis revealed that *ImCoh* was selectively modulated along cross-hemispheric diagonal pathways rather than within same-hemisphere pairs **(Figure 4A)**. Specifically, we observed a significant enhancement in 𝛽 band synchronization between the contralateral prefrontal cortex (relative to the target) and the ipsilateral visual cortex compared to contralateral visual cortex **(Figure 4B)**. This increase in *ImCoh* emerged within the stimulus epoch (∼0.15s) and occurred in several discrete intervals during the delay period, characterized by several significant temporal points (*p* < 0.05, not corrected for multiple comparisons). On the other hand, a decoupling was observed between ipsilateral prefrontal cortex and contralateral visual cortex compared to ipsilateral visual cortex **(Figure 4C)**, starting after stimulus epoch and occurring in several discrete intervals throughout the delay.

**Figure 4:**
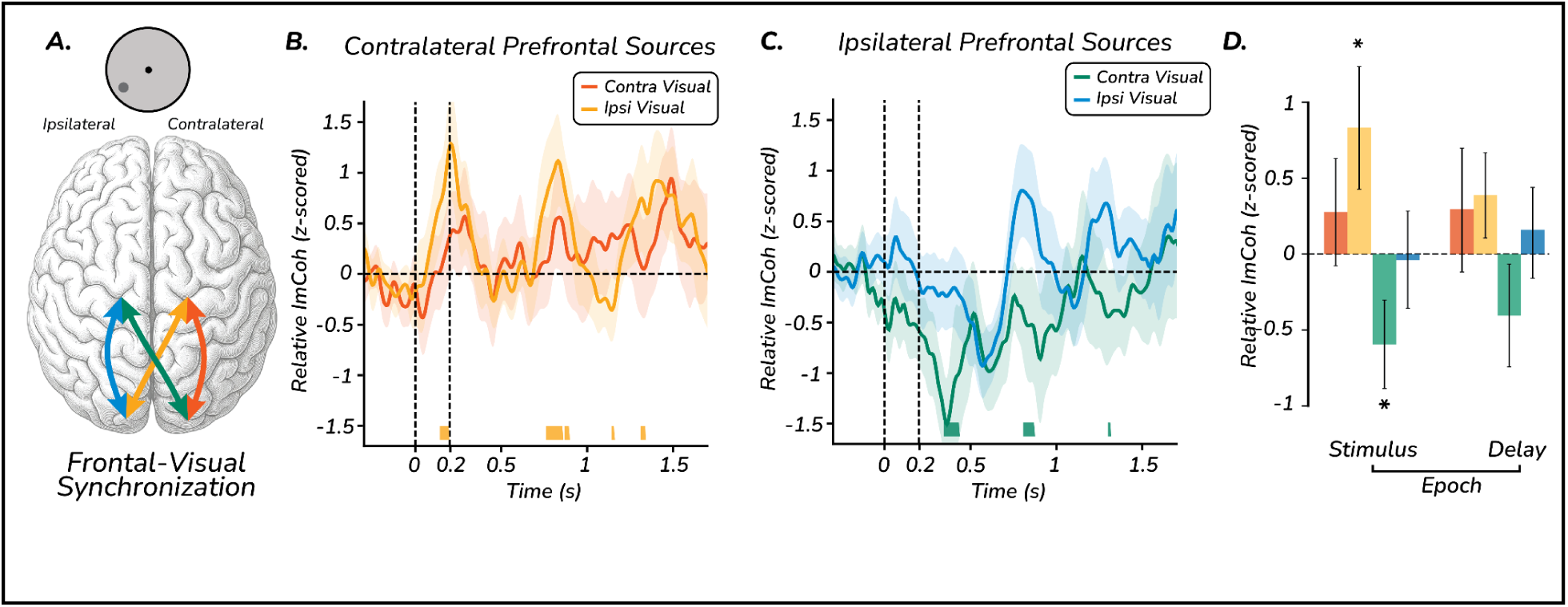
Cross-hemispheric. 𝛽 **band synchronization between prefrontal and visual cortex. A.** Schematic of 𝛽 band imaginary coherence (ImCoh) analysis. Arrows represent functional connectivity between prefrontal and visual regions, color-coded by their hemispheric relationship to the target stimulus (depicted here for target present in left visual field, gray dot). Orange and green arrows indicate cross-hemispheric synchrony, while red and blue arrows indicate same-hemispheric synchrony. **B, C.** Time-resolved relative ImCoh (z-scored) compared to fixation baseline. (B) Contralateral prefrontal sources (relative to target) exhibit significant synchronization with the ipsilateral visual cortex (orange line) compared to the contralateral visual cortex (red line). (C) Ipsilateral prefrontal sources exhibit significant suppression of synchronization (decoupling) with contralateral visual cortex (green line) relative to ipsilateral visual cortex (blue line). Shaded areas represent SEM, and horizontal bars on the x-axis indicate significant temporal points (*p* < 0.05, not corrected for multiple comparisons). **D.** Statistical summary of relative ImCoh changes across Stimulus (0.1 - 0.3s) and Delay (0.6 - 1.5s) epochs. A significant interaction is observed specifically along cross-hemispheric diagonal pathways during the stimulus epoch: contralateral prefrontal sources show enhanced coupling with ipsilateral visual cortex (orange), while ipsilateral prefrontal sources show decoupling from contralateral visual cortex (green). Same-hemispheric connectivity (red and blue bars) remains at baseline levels throughout the task. Asterisks indicate significant deviations from baseline (*p* < 0.05). Error bars represent SEM.

To quantify these effects across the task, we compared the average *ImCoh* changes during the Stimulus (0.1–0.3s) and Delay (0.6–1.5s) epochs **(Figure 4D)**. During the stimulus epoch, we observed an enhanced synchronization between contralateral prefrontal and ipsilateral visual cortices (*z*-scored *ImCoh*: 0.833 ± 0.403, *p_perm* = 0.047, *d* = 0.45) and a decoupling between ipsilateral prefrontal and contralateral visual cortices (*z*-scored *ImCoh*: -0.596 ± 0.290, *p_perm* = 0.048, *d* = -0.45). Synchrony within the same hemisphere pairs remained at baseline levels for both contralateral (*z*-scored *ImCoh*: 0.278 ± 0.353, *p_perm* = 0.45, *d* = 0.17) and ipsilateral (*z*-scored *ImCoh*: -0.040 ± 0.316, *p_perm* = 0.90, *d* = -0.03) hemispheres.

Similarly, during the delay epoch (0.6 - 1.5s), the synchronization between contralateral prefrontal and ipsilateral visual cortices (*z*-scored *ImCoh*: 0.388 ± 0.279, *p_perm* = 0.18, *d* = 0.30) remained elevated but did not reach significance, and the ipsilateral prefrontal and contralateral visual cortices remained decoupled (*z*-scored *ImCoh*: -0.406 ± 0.345, *p_perm* = 0.25, *d* = -0.26) but did not reach significance. Within hemisphere synchronization during the delay also remained near baseline for both contralateral (*z*-scored *ImCoh*: 0.297 ± 0.401, *p_perm* = 0.47, *d* = 0.16) and ipsilateral (*z*-scored *ImCoh*: 0.160 ± 0.279, *p_perm* = 0.58, *d* = 0.13) hemispheres.

This provides evidence that prefrontal cortex and visual cortex exhibit a cross-hemispheric synchrony, with the prefrontal cortex likely providing an inhibitory scaffolding pattern over visual cortex, thereby retaining high-fidelity memory contents in visual cortex.

## Discussion

We leveraged the high temporal resolution of MEG to identify the neural coordination supporting human visuospatial working memory. We report three key findings: 1) 𝛽 band power persisted throughout the working memory delay in visual and prefrontal cortical regions, 2) 𝛽 band power in visual cortex, but not prefrontal cortex, tracked working memory content and predicted memory performance, and 3) memory maintenance is characterized by a discrete, cross-hemispheric 𝛽 band synchrony between prefrontal and visual cortices.

We observed enhanced and persistent 𝛽 band activity in visual and prefrontal cortices during working memory delay. Moreover, in visual cortex, during the delay, the topography of 𝛽 band power varied continuously as a function of stimulus location. Specifically, relative to the spatial position of the target, we observed a desynchronization of 𝛽 band power in contralateral hemisphere, the hemisphere involved in processing and maintaining the stimulus location in working memory (Pfurtscheller and da Silva, 1999; Klimesch, 2012). At the same time, we observed synchronized 𝛽 band power in ipsilateral hemisphere, the hemisphere that is not involved in stimulus processing. This contralateral desynchronization is a well-established signature of spatial working memory and has been observed in both EEG and MEG studies (Medendorp et al., 2007; Van Der Werf et al., 2008; van Dijk et al., 2010). Importantly, these oscillatory shifts do not merely mirror passive visual processing, but actively encode the coordinates of the spatial motor goals preceding memory-guided saccades <u>(Van Der Werf et al. 2008)</u>. The topography of lower frequency neural activity in α/𝛽 power reflects local inhibition, serving to suppress the task-irrelevant visual field in order to prevent interference with the stored memory trace (Worden et al., 2000; Zumer et al., 2014). Large populations synchronize spiking in 𝛽-band when the neural activity of the population is inhibited (Lundqvist et al., 2016; Miller et al., 2018), effectively gating sensory processing by reducing the excitability of local neuronal populations. A reliable way to sculpt information in sensory cortex, which has constant excitatory feedforward input, is to pattern it with an inhibitory signal to suppress sensitivity to irrelevant information (Jensen and Mazaheri, 2010).

This topography of 𝛽-band activity can either track precise spatial location (Foster et al., 2017) in memory or simply reflect a lateralized control signal. We show that this topographic change of 𝛽-band power tracked the precise stimulus location at a trial-by-trial level in visual but not prefrontal cortex, reflecting that topography of neural activity in 𝛽-band tracks memory content. This suggests a strong functional coupling between the pattern of neural activity and the readout used for memory-guided behavior. The presence of memory contents in neural activity in visual but not prefrontal cortex is consistent with previous findings in human fMRI studies (Harrison and Tong, 2009; Serences et al., 2009; Rahmati et al., 2018; Kwak and Curtis, 2022; Duan and Curtis, 2024). Our results provide mechanistic support for the sensory recruitment model, which posits that association cortices modulate sensory areas to retain high-fidelity representations (Curtis and D’Esposito, 2003; Postle, 2006; Serences et al., 2009). An important observation to note here is that, unlike fMRI studies where an increased BOLD amplitude is used to decode memory content in early visual cortex (Harrison and Tong, 2009; Serences et al., 2009; Pratte and Tong, 2014; Ester et al., 2015), we show the existence of a complementary signal with increases in non-stimulated hemifield in 𝛽-band activity tracked memory information as well. Indeed, the local spiking in early sensory areas is modulated by local 𝛽 activity (Lundqvist et al., 2016) in a complementary fashion, with increases in power in 𝛽-band activity associated with decreases in local spiking.

The existence of 𝛽-band activity in both prefrontal and visual cortices during working memory delay suggests that the two regions are likely communicating via establishing a common synchronized channel (Helfrich and Knight, 2016; Comeaux et al., 2023). We investigated this using imaginary coherence in 𝛽-band between prefrontal and visual cortices. We observed that contralateral prefrontal cortex exhibits an enhanced synchronization with ipsilateral visual cortex, the visual hemisphere with high 𝛽-band power. At the same time, the ipsilateral prefrontal cortex decoupled from contralateral visual cortex. This suggests a cross-hemispheric interaction between prefrontal and visual cortices, with enhanced synchrony with visual hemisphere inhibited during retention and decreased synchrony with the hemisphere active, suggesting prefrontal control being largely inhibitory in nature (Bastos et al., 2015). This synchronized activity was not constant throughout the delay, but appeared to be dynamic, suggesting that the two areas might be communicating via a recurrent loop to keep the contents of memory intact in visual cortex. Prefrontal cortex likely acts as a neural index or pointer (Kriete et al., 2013) rather than storing literal sensory features of the memorandum. This implies that the prefrontal control is only as effective as the fidelity of representation it recruits in visual cortex (Postle, 2006), effectively decoupling executive control (the memory address) from physical storage (the memory content).

A significant anatomical question raised by our findings is the pathway supporting the cross-hemispheric synchronization between prefrontal and visual cortices **(Figure 4A)**. However, direct monosynaptic projections across these distal nodes are absent. This likely necessitates the role of an intermediate relay. We propose that the mediodorsal thalamus (Halassa and Kastner, 2017) or the pulvinar (Saalmann et al., 2012) may act as a central router to mediate these interactions. The mediodorsal thalamus is known to sustain cortical delay activity and possesses the requisite anatomical connectivity to synchronize disparate cortical regions into a unified functional network (Schmitt et al., 2017). Indeed, recruitment through coherence hypothesis suggests a potential role of thalamus in generating the synchrony between prefrontal and relevant sensory areas, which is required to gate information in working memory (Comeaux et al., 2023). Further studies investigating the role of thalamocortical loops and 𝛽 dynamics will be essential in understanding how the cross-hemispheric synchronization of the prefrontal-visual network is physically instantiated.

In conclusion, our results demonstrate that working memory is neither a passive storage process in sensory cortex nor an isolated executive function in prefrontal cortex, but rather a dynamic coordination of the two, and likely several other distributed modes of the network (Christophel et al., 2017). By leveraging 𝛽-band synchronization, prefrontal cortex functions as an executive controller, recruiting and patterning sensory populations to transform transient sensory inputs into stable internal representations.

## Acknowledgements

The work was supported by NIH R01s EY033925 and EY016407 to CEC. We thank Sebastian Michelmann for analytic support.

## Author Contributions

C.E.C. and S.D. conceived and designed research; S.D. performed experiments; M.D. analyzed data and prepared figures; M.D., S.D., and C.E.C. interpreted results of experiments; M.D. and C.E.C. drafted manuscript; M.D. and C.E.C. edited, revised, and approved final version of manuscript.

## Data Availability Statement

The raw and preprocessed magnetoencephalography (MEG) data, behavioral data, and custom MATLAB/Python analysis scripts used in this study are hosted on the Open Science Framework (OSF) and can be accessed at https://osf.io/653sn or via the corresponding author.

