## Supplementary Figures for "Neural synchrony between prefrontal and visual cortex supports visual working memory"

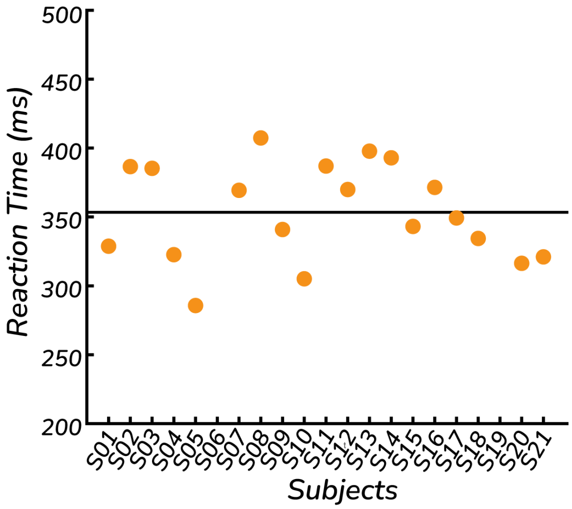


**Supplementary Figure 1: Saccade Reaction time for memory-guided saccade task**
Mean saccade reaction time (ms) across participants (*N* = 21). The horizontal line indicates group mean error 356 ± 8 ms.

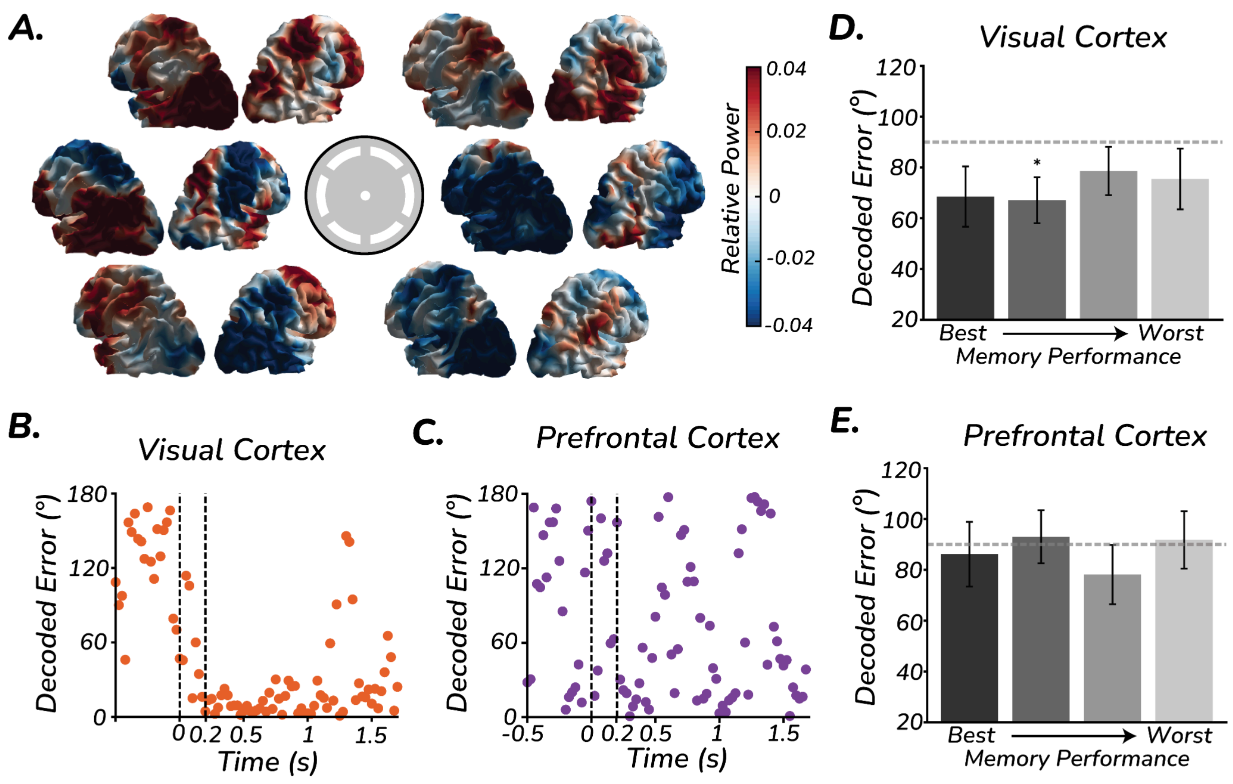


**Supplementary Figure 2: Visual α-band activity tracks target location but does not predict memory performance.**

**A.** Topography of α-band power over the cortical sheet in MNI space during the working memory delay, binned into six spatial target locations (wedges in the center). Relative power was calculated as the power at a specific target bin location relative to the mean power across all other bins. Note the characteristic spatial profile: enhanced α-band power in the ipsilateral hemisphere and suppressed power in the contralateral hemisphere relative to the target, with a rotational topography that shifts as a function of target polar angle.

**B, C.** Temporal dynamics of spatial decoding. Decoding error (degrees) for target location over time from α-band power in visual (B, orange) and prefrontal (C, purple) ROIs. Vertical dashed lines indicate stimulus onset and offset. Target location is reliably decoded from visual sources throughout the delay, whereas decoding from prefrontal sources remains at chance levels.D, E. Neural decoding does not predict working memory performance. Trials were divided into four quantiles (Q1–Q4) based on memory error (best to worst, left to right). In contrast to $\beta$-band dynamics, decoding error in the visual cortex (D) did not significantly correlate with behavioral performance, with decoding precision remaining stable across all performance quantiles. Similarly, no relationship between behavior and decoding was observed in the prefrontal cortex (E). Error bars represent SEM; * denotes significant difference from chance (90°, $p < 0.05$).
